# Visualizing cellular and tissue ultrastructure using Ten-fold Robust Expansion Microscopy (TREx)

**DOI:** 10.1101/2021.02.03.428837

**Authors:** Hugo G.J. Damstra, Boaz Mohar, Mark Eddison, Anna Akhmanova, Lukas C. Kapitein, Paul W. Tillberg

## Abstract

Expansion microscopy (ExM) is a powerful technique to overcome the diffraction limit of light microscopy that can be applied in both tissues and cells. In ExM, samples are embedded in a swellable polymer gel to physically expand the sample and isotropically increase resolution in x, y and z. The maximum resolution increase is limited by the expansion factor of the gel, which is four-fold for the original ExM protocol. Variations on the original ExM method have been reported that allow for greater expansion factors but at the cost of ease of adoption or versatility. Here, we systematically explore the ExM recipe space and present a novel method termed Ten-fold Robust Expansion Microscopy (TREx) that, like the original ExM method, requires no specialized equipment or procedures. We demonstrate that TREx gels expand ten-fold, can be handled easily, and can be applied to both thick mouse brain tissue sections and cultured human cells enabling high-resolution subcellular imaging with a single expansion step. Furthermore, we show that TREx can provide ultrastructural context to subcellular protein localization by combining antibody-stained samples with off-the-shelf small molecule stains for both total protein and membranes.

## INTRODUCTION

Expansion Microscopy (ExM) circumvents the diffraction limit of light microscopy by physically expanding the specimen four-fold in each dimension (F. Chen, Tillberg, & Boyden, 2015; Tillberg et al., 2016). Expansion is achieved by chemically anchoring proteins and other biomolecules directly to a hyper-swelling gel, followed by aggressive proteolysis to enable uniform swelling of the gel material. While other super-resolution approaches are not readily compatible with thick tissue slices and require specialized optics (Hell & Wichmann, 1994), fluorophores (Rust, Bates, & Zhuang, 2006), or software (Gustafsson, 2000), ExM is compatible with any microscope (R. Gao et al., 2019; Zhang et al., 2016), including other super resolution modalities (M. Gao et al., 2018; Halpern, Alas, Chozinski, Paredez, & Vaughan, 2017; Xu et al., 2019), and performs well in both cultured cells and thick tissue slices (F. Chen et al., 2015; Tillberg et al., 2016). Assuming sufficiently high labeling density, the resolution increase of ExM depends on the expansion factor of the gel recipe used. Recently, ExM variants have been described that seek to improve resolution by increasing the expansion factor. For example, iterative ExM (iExM) uses sequential embedding in multiple expansion gels to achieve 15x and greater expansion but requires a complex sequence of gel re-embedding, link cleaving, and fluorophore transfer (Chang et al., 2017), limiting its broad adoption.

The expansion factor of the gel itself can be improved by decreasing the concentration of crosslinker (Okay, 2009), usually bisacrylamide (bis), although this is generally at the expense of the mechanical integrity of the gel. For example, reducing the bis concentration in the original ExM recipe from 1.5 to 0.25 ppt (parts per thousand) produces a ~9-fold expanding gel (Chen, Tillberg and Boyden, 2015, SF5), but these gels are too soft to hold their shape under the force of gravity. As a result, they are difficult to handle without breaking and display non-uniform expansion. This tradeoff of expansion versus gel mechanical integrity has not been explored in a quantitative or systematic way.

Another gel recipe variant, using a high concentration of the monomer dimethylacrylamide (DMAA), has enough crosslinking through side reactions and polymer chain entanglement that the crosslinker can be omitted entirely, producing ~10-fold expansion in one step (Truckenbrodt et al., 2018). This recipe has been used to expand cultured cells and thin cryosectioned tissue (Truckenbrodt, Sommer, Rizzoli, & Danzl, 2019), but reportedly requires rigorous degassing to remove oxygen prior to gelation, making it laborious to use. Moreover, expansion of thick tissue slices (>50 μm) has not been demonstrated using this method. Thus, a robustly validated and easily adoptable method that is compatible with multiple sample types and enables single step expansion well over 4x without compromising gel integrity is lacking.

Here, we explored the expansion gel recipe space in a systematic manner, assessing the limits of single-round expansion using reagents and methods that would be familiar to labs already performing ExM. For any given choice of recipe parameters (monomer concentrations, gelation temperature, initiator concentration, etc.), varying the crosslinker alone yielded a family of recipes whose expansion factor and mechanical quality vary smoothly from high expanding, mechanically unstable to low expanding, tough gels. A range of crosslinker concentrations was tested for each family because the optimal crosslinker concentration may vary by family. From this exploration we generated TREx, an optimized ExM method that allows for robust ten-fold expansion in a single step. We show TREx can be used to expand thick tissue slices and adherent cells, and is compatible with antibodies and off-the-shelf small molecule stains for total protein and membranes. Together, we show that TREx enables 3D nanoscopic imaging of specific structures stained with antibodies in combination with cellular ultrastructure.

## RESULTS

To systematically explore the expansion recipe space, we developed a streamlined approach for synthesizing dozens of gel recipes and characterizing their mechanical quality in parallel. For every set of gel recipe parameters (component concentrations and gelation temperature, listed in Fig. 1A) we define a recipe family as the set of recipes generated by varying the crosslinker (bisacrylamide) concentration. For each family, we tested five recipes with crosslinker concentrations log-spaced from 1000 to 10 ppm (parts per million, or μg/mL), plus one with zero crosslinker. We also included the original ExM recipe, with 0.15% (1500 ppm) crosslinker. For each recipe, we cast three gel specimens, expanded them fully in water, and measured the expansion factor (Fig. 1B). We found that resistance to deformation under the force of gravity was a good proxy for the more subjective judgement of ease of gel handling. We measured gel deformation by placing a semicircular punch from each gel upright on its curved edge and allowing the gel corners to deflect under the force of gravity. We defined the deformation index as the vertical displacement of each gel corner, divided by the gel radius (Fig. 1C), which ranges from 0 (for gels that do not deform) to 1 (for gels that deform freely under their own weight). We manually calibrated this measurement, finding that deformation indices between 0 and 0.125 corresponded to gels with excellent ease of handling, 0.125 to 0.25 corresponded to acceptable ease of handling, and anything higher than 0.25 was unacceptable. While not as theoretically informative as elastic modulus and yield strength measurements, this measurement can be repeated and extended to new gel recipes by any lab developing expansion methods, without access to specialized equipment. We plotted the deformation index for each gel as a function of its expansion factor (Fig. 1D) to directly assess the tradeoff between expansion and mechanical quality.

**FIGURE 1:**
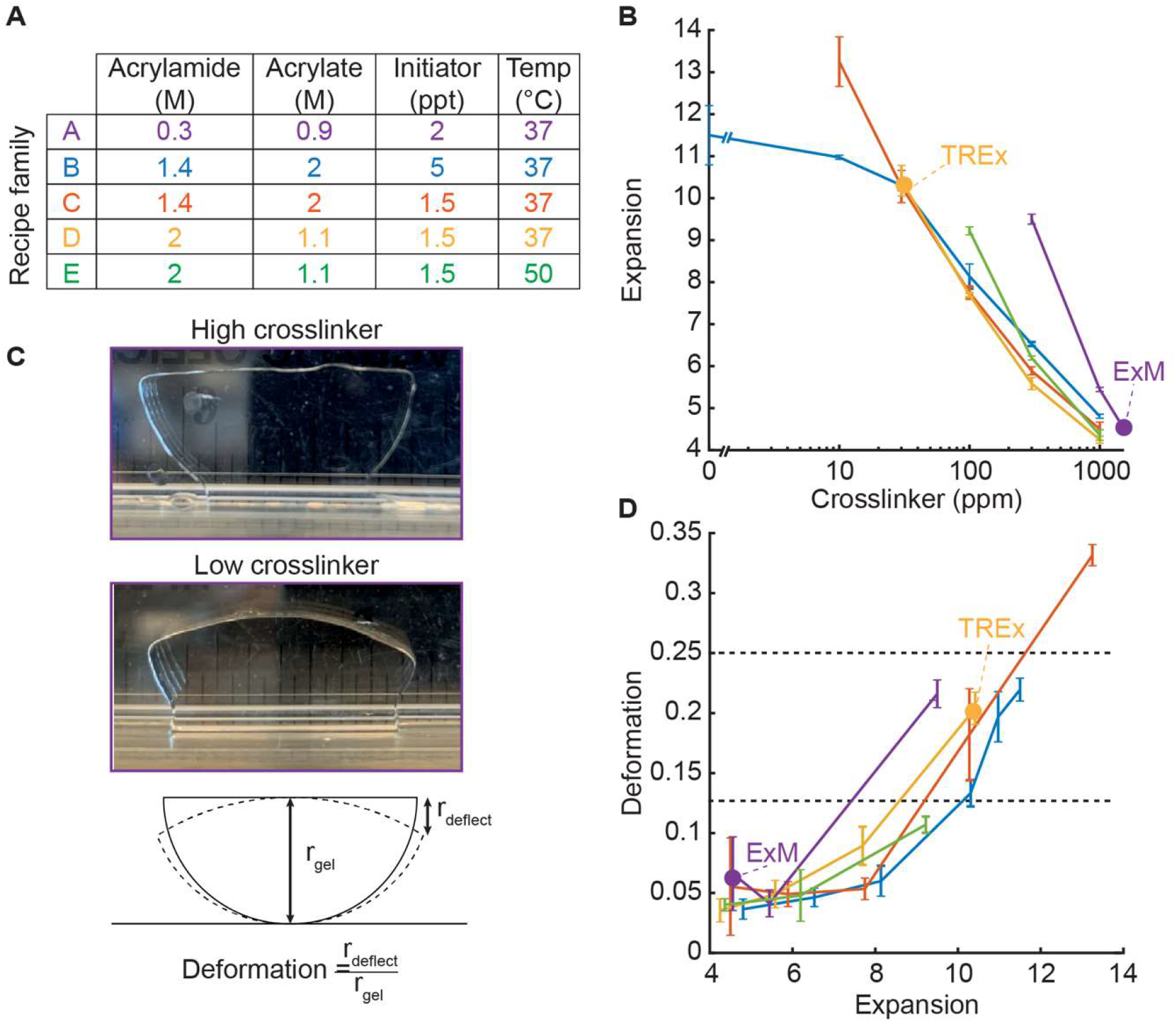
Development of TREx gel recipe. A) Parameters of gel recipe families explored, including component concentrations and gelation temperature. Each family was characterized by keeping these conditions constant while systematically varying the crosslinker concentration. B) Expansion factor (mean ± S.D., n=3) versus crosslinker concentration (log scale) for each gel recipe family without biological specimens. Line colors correspond to recipe families as in 1A. Specific recipes are indicated with a filled purple dot (original ExM recipe) and yellow dot (TREx). All recipe families were tested with crosslinker concentrations of 0, 10, 30, 100, 300, 1000 ppm, plus an additional condition for family A with 1500 ppm, corresponding to the original ExM recipe. Only conditions in which gels formed are plotted. C) Definition of gel deformation index. Example gels from recipe family A with high crosslinker and low deformation (top panel, 1.5 ppt), and low crosslinker and high deformation (middle panel, 300 ppm). Bottom panel, schematic illustrating deformation index measurement. D) Deformation index (mean ± S.D., n=3) versus expansion factor for each gel recipe family without biological specimens, with line colors and dots corresponding to specific recipes as in 1A and 1B. Horizontal grey lines indicate thresholds for gels with mechanical quality deemed perfect (deformation < 0.125) and acceptable (deformation < 0.25). Ideal recipes would occupy the lower right quadrant, corresponding to high expansion and low deformability.

### Development of the TRex gel recipe

We began by characterizing a recipe family generated from the original ExM recipe (family A). Consistent with (F. Chen et al., 2015), reducing the crosslinker to 300 ppm increased the expansion to ~9x, below which the gels fail to form consistently (Fig. 1B, purple). We next characterized a high-monomer recipe family (family B) inspired by the 4x-expanding Ultra-ExM recipe (Gambarotto et al., 2019). Gambarotto et al. found that a higher monomer concentration relative to the original ExM recipe was necessary for high fidelity preservation of the shape of centrioles. This was offset with a lower crosslinker (bisacrylamide) concentration of 0.1% (1000 ppm) to achieve 4x expansion. Indeed, for this high-monomer family of recipes, expansion as a function of crosslinker concentration was shifted leftward compared to standard ExM (Fig. 1B, blue). As the crosslinker was decreased below 30 ppm, the increase in expansion factor saturated around 11.5x. The deformation index versus expansion factor curve for the high monomer family ran below that for standard ExM, indicating that for a given expansion factor the high-monomer gel holds its shape better than the corresponding standard ExM gel (Fig. 1D, blue).

Compared with standard ExM, this high monomer family uses a higher concentration of radical initiator and accelerator to trigger polymerization (5 ppt each of APS and TEMED, versus 2 ppt in standard ExM). We found that this high initiation rate causes gels to form within minutes at room temperature. Because the rates of initiation and polymerization increase with temperature, it is likely that specimens are not fully equilibrating to the gelation temperature of 37 °C before the onset of gelation, introducing a potential source of experimental variability. This rapid gelation makes the gelation chamber assembly step more time sensitive and presents challenges for adapting the technique to thick tissue slices, as thick tissue slices require extra time for the gelation solution to diffuse throughout the sample prior to polymerization. Therefore, we also tested the same high monomer recipe family but with initiator and accelerator reduced to 1.5 ppt (family C). The expansion versus crosslinker curve for this family was similar to family B for high crosslinker concentrations but displayed a slightly greater slope. Unlike family B, the expansion factor did not saturate upon decreasing the crosslinker concentration. Instead, the expansion factor continued to increase to 13x expansion at 10 ppm crosslinker (Fig. 1B, red), with zero-crosslinker gels failing to form. Family C enables ten-fold expansion without sacrificing acceptable gel mechanical quality (30 ppm bis, Fig. 1D, red), and without the faster, less controlled gelation kinetics of family B.

The recipe families explored above (B, C) feature a high fraction of sodium acrylate relative to acrylamide. Acrylate drives expansion of the gel but comes in widely varying purity levels and, in some cases, causes tissue to shrink. This macroscopic tissue shrinkage is modest compared to the gel expansion but may not be uniform at all scales. We therefore tested an alternative recipe family (D) with higher acrylamide to acrylate ratio (2.1:1). Increasing the acrylamide to acrylate ratio did not change the expansion factors appreciably at a given crosslinker concentration, suggesting that the swelling effect of acrylate saturates at high concentrations. At the maximum expansion factor of ~10, the deformation behavior was comparable to family C. We chose to proceed with family D due to its lower acrylate content.

We further tested an elevated gelation temperature of 50 °C (family E), in an attempt to increase the initiation rate without introducing premature gelation as seen in recipe family B. Compared to family D, the expansion factors were around 15% higher at 100 ppm (6x) and 300 ppm crosslinker (9x), but gelation failed at lower concentrations, leaving family D as the family with a higher maximum expansion factor (i.e., 10x at 30 ppm bis). The deformation versus expansion curve for family E was similar to the other high monomer recipe families (Fig. 1D, green), but was found to be sensitive to processing details, such as the gelation chamber construction and placement within the incubator. This suggests that premature gelation prior to equilibrating at the higher temperature reduces the replicability of this recipe family.

Considering all 5 recipe families, family B (high acrylate and high APS/TEMED) displayed the lowest deformation index for a given expansion factor. Family D (high acrylamide and low APS/TEMED) displayed similar performance, with the deformation index remaining well within the acceptable range for expansion factors up to 10. In handling high-expanding (>8x) gels from all recipe families, we found that while those from the standard ExM family (A) were extremely prone to fragmentation, those from any of the high monomer families could be handled more easily (and even dropped from a height of several feet) without breaking. Because the reduced initiator concentration of family D results in a slower and more controlled polymerization rate, and because we preferred a lower acrylate content, we chose this recipe family to proceed to biological specimen expansion. We found that the exact expansion factor varied for different specimens and gelation chamber geometries but could readily be adjusted by fine-tuning the crosslinker concentration. We thus recommend that users test gels with a range of crosslinker concentrations between 30 and 100 ppm to find a suitable recipe for their specimen preparation. We name the resulting method Ten-fold Robust Expansion (TREx) microscopy.

### Subcellular imaging of specific proteins and cellular ultrastructure in thick brain slices

In electron microscopy, non-specific stains for proteins and membranes are commonly used to provide structural detail at high spatial resolution. Recently, the use of non-specific NHS ester protein stains and other small molecule probes has been combined with ExM (M’Saad & Bewersdorf, 2020; Mao et al., 2020; Yu et al., 2020). Expansion allows visualization of intracellular detail in such densely stained samples, which would otherwise be too crowded to lead to meaningful contrast. These applications have the promise to bring together the advantages of light microscopy (specific staining using antibodies and volumetric imaging) with the advantage of seeing cellular context typically provided by electron microscopy. Because TREx reaches single-step expansion factors at which small-molecule stains are expected to be useful, we set out to explore this idea further.

We applied BODIPY-FL NHS dye (total protein stain; see Materials and Methods) after expansion with TREx to demonstrate total protein distribution in thick (100 μm) slices of mouse brain cortex (Fig. 2A, Fig. 2—fig. supp. 1, and Fig. 2—supp. movie 1). The neuropil region outside the cell somas contained a rich profusion of fibers and structures visible in sharp relief. The nucleus of each cell was easily identified, with especially strong staining in nucleoli-like structures. Surrounding each nucleus, the nuclear envelope could be identified, with particularly dense total protein stain on the side facing the nucleus. The nuclear envelope was punctuated by heavily stained spots that span the envelope, consistent with nuclear pore complexes (Fig. 2A, inset). Within the cytosol, several organelles were marked by either heavy inner staining with a dim border or weak inner staining.

**FIGURE 2:**
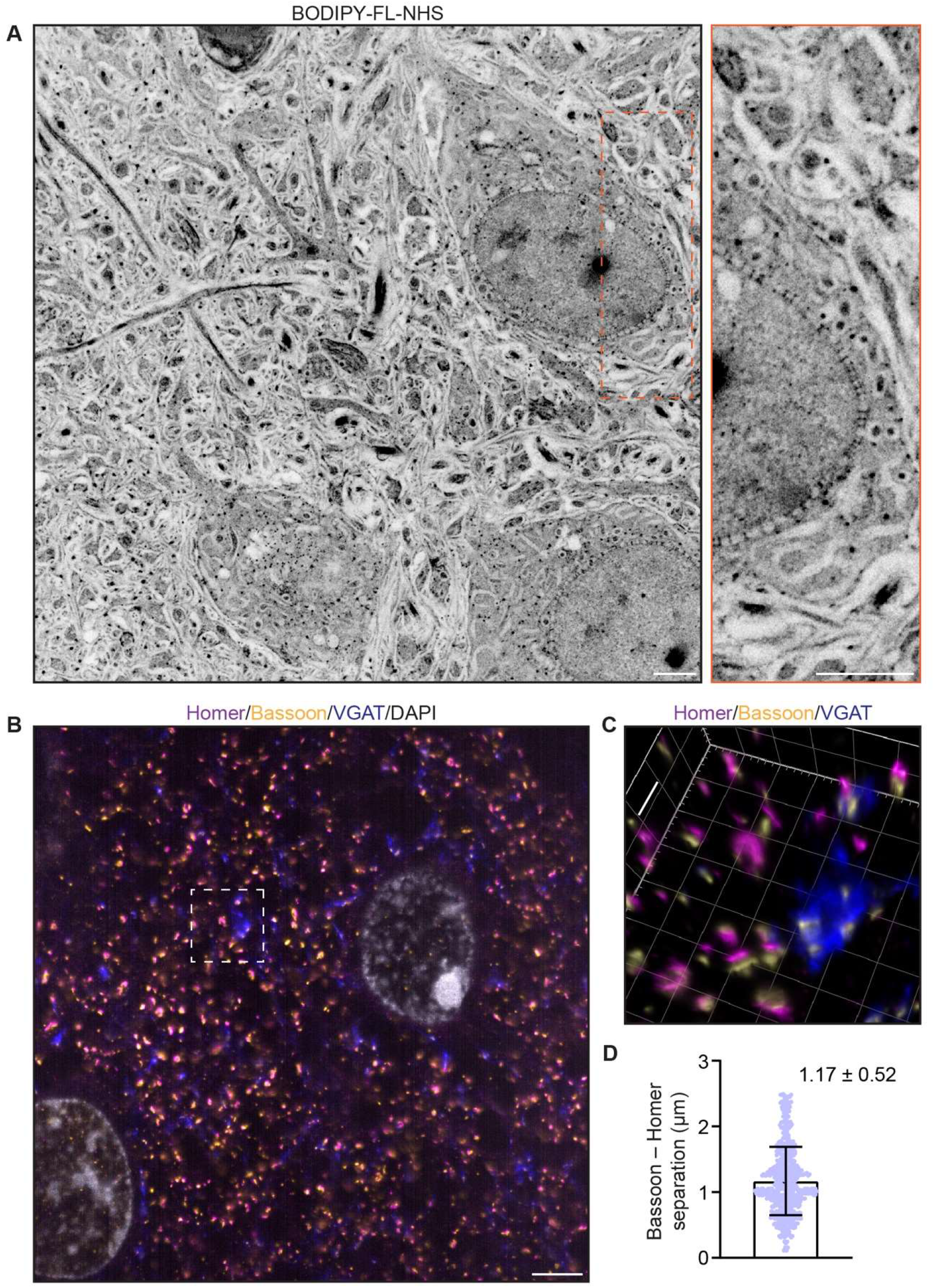
TREx in mouse brain tissue slices. A) Mouse brain tissue (cortex) expanded using TREx, stained for total protein content with BODPIY-FL NHS, and imaged by confocal microscopy. Displayed contrast is inverted to show dense stained regions as dark. Inset, zoom-in showing nuclear envelope with densely stained structures spanning the nuclear envelope, consistent with nuclear pore complexes. B) Mouse brain tissue (cortex) stained with antibodies against homer (magenta), bassoon (yellow), and VGAT (blue), and DAPI (grey), and expanded using TREx. C) Volumetrically rendered zoom-in of white box in (A) showing paired Bassoon- and Homer-rich structures, consistent with excitatory synapses. Depending on the orientation, clear separation of Bassoon and Homer can be observed, as well as a complex, structured pre-synaptic vesicle pool marked by VGAT bearing several release sites marked by Bassoon. D) Quantification of Bassoon and Homer separation (mean ± S.D. plotted, n=538 synapses, 1 replicate). Scale bars (corrected to indicate pre-expansion dimensions): main ~2 μm, zooms ~400 nm

We attempted to optimize protein retention, according to the total protein stain intensity, by reducing both protein anchoring and proteolysis compared with the original ExM. We tested a range of anchoring strengths by varying the concentration of the acryloyl-X SE (AcX) anchoring molecule applied prior to gelation. This was done in combination with two reduced disruption methods: proteinase K applied at one tenth that of the original ExM method (Fig. 2—fig. supp. 1, top row), and a high-temperature, protease-free denaturation treatment (Gambarotto et al., 2019; Ku et al., 2016; Tillberg et al., 2016; Zwettler et al., 2020) similar to that employed in Western blotting (Fig. 2—fig. sup. 1, bottom row). The protease-free treatment enabled greater protein retention but at the cost of incomplete expansion. This could be offset through reduced AcX concentration, though this was not clearly superior to high AcX followed by proteolysis, indicating a general tradeoff between protein retention and gel expansion (see Materials and Methods). We chose a hybrid approach with moderate AcX anchoring and low concentration proteinase K digestion followed by high temperature denaturation to proceed.

We next tested whether antibodies, applied to the tissue using a standard immunofluorescence procedure before embedding, were also retained in the TREx gel. We stained mouse brain cortex tissue for Bassoon (a marker for both excitatory and inhibitory pre-synaptic active zones), Homer (a marker for the excitatory post-synaptic apparatus), and VGAT (a vesicular GABA transporter in the pre-synaptic compartments of inhibitory synapses). After staining and anchoring with AcX, tissue was expanded with TREx and imaged by light sheet microscopy. Numerous putative excitatory synapses were observed at high density, with clearly separated Bassoon and Homer pre- and post-synaptic staining (Fig. 2B-C and Fig. 2— supp. movie 2). Because of the excellent axial resolution, TREx allowed us to quantify the separation of Bassoon and Homer regardless of the angle of the synapse with respect to the imaging plane (Fig. 2D). We found an average separation of 1.17 μm ± 0.52 μm (mean ± S.D.,583 synapses), which, when corrected for expansion, is consistent with previous reports in cultured neurons that estimated the synapse separation between 90-130 nm (Glebov, Cox, Humphreys, & Burrone, 2016; Wiesner et al., 2020). Compared with Bassoon and Homer, VGAT had a more extended staining pattern, consistent with the known distribution of synaptic vesicles throughout pre-synaptic boutons. Elaborately shaped compartments with dense VGAT staining were seen with multiple synaptic release sites marked by Bassoon (Fig. 2C). As expected, these release sites were not paired with the excitatory post-synaptic marker Homer. These results demonstrate the ability of TREx to preserve correct synaptic staining while enabling sub-diffraction limited imaging of large tissue sections.

### Validation of expansion factor and deformation

Increasing the expansion factor from 4 to 10x could result in greater sensitivity of the expansion factor to local variation, for example in protein dense complexes, resulting in less uniform expansion. To examine this, we explored the nanoscale isotropy of TREx by imaging nuclear pore complexes (NPCs), which have a highly stereotyped and well characterized structure on the sub-100 nm scale. NPCs have recently been explored as a reference structure for super-resolution microscopy methods, including ExM in combination with other super-resolution methods (Pesce, Cozzolino, Lanzanò, Diaspro, & Bianchini, 2019; Thevathasan et al., 2019). For the conventional 4-5x expansion approach, this revealed that the diameter of the NPC was 14-29% smaller than expected from the macroscopic expansion of the gel. We used a NUP96-GFP homozygous knock-in cell line (Thevathasan et al., 2019) to study the quality of nuclear pore expansion using TREx with well validated anti-GFP antibodies (Fig. 3A). After expansion with TREx, individual NPCs were uniformly retained and clearly visible using diffraction-limited confocal microscopy (Fig. 3B). An antibody against NUP153 similarly demonstrated individual NPCs but with less complete NPC coverage compared with the antibody stain against the NUP96-GFP tag (Fig. 3C). The macroscopic gel expansion factor was 9.5x, suggesting an expected NPC size after expansion of 107 nm x 9.5 = 1.02 μm. We used a semi-automated approach to determining the diameter of 60 NPCs randomly chosen from three non-adjacent cells and found the size after expansion to be 939 nm ± 90 nm (mean ± S.D.) (Fig. 3D). This is about 8% smaller than expected based on the macroscopic expansion of the gel and implies a local expansion factor of 8.8x, or 92% of the expected 9.5x. These data indicate that TREx offers more uniform local expansion compared to conventional ExM.

**FIGURE 3:**
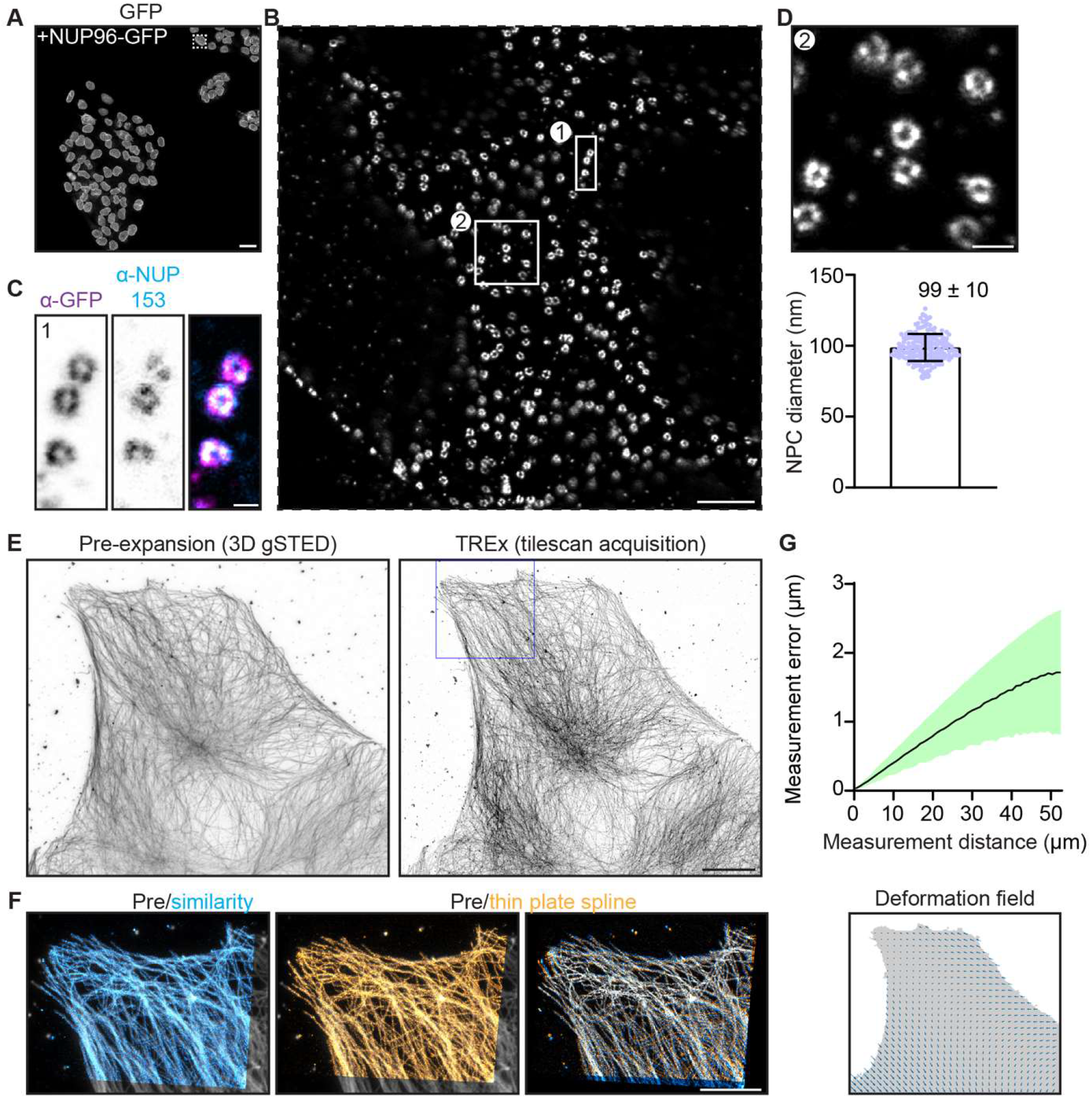
Characterization of expansion isotropy using TREx. A) U2OS knock-in cells with homozygous NUP96-GFP, amplified with anti-GFP antibodies. B) One nucleus from boxed region of (A), imaged by confocal microscopy after TREx. C) High-resolution view of several nuclear pores from boxed region (1) of panel (B), showing both anti-GFP (magenta) and anti-NUP153 (endogenous nuclear pore protein, cyan) staining. D) High-resolution view of several nuclear pores from boxed region (2) of panel (B) (top). Distribution of diameters of individual nuclear pores (bottom), corrected for the macroscopic expansion factor of 9.5x. N=60 NPCs from 3 spatially separated cells. E) Maximum projection of pre-expansion 3D gSTED acquisition (left) and maximum projection of tilescan acquisition (42 tiles, post expansion size ~750×650 um) of the same cell post-expansion (right). F) Post expansion single field of view, as indicated with magenta box in (D), aligned with the pre-expansion image (grey) by similarity transformation (cyan) or thin plate spline elastic transformation (orange). Right shows overlay of similarity and elastic transformation to illustrate local deformations. G) Quantification of measurement errors of the stitched dataset due to non-uniform expansion. Mean error for a given measurement length (black line) ± S.D. (shaded region). The residual elastic deformation field is shown below. Scale bars: A ~1 μm, B ~50 μm, C ~100 nm, D ~200 nm, E ~10 μm, F ~5 μm.

We further quantified the measurement error introduced by non-uniform expansion by comparing antibody-stained microtubules imaged before expansion with 3D gSTED versus after expansion with confocal microscopy (Fig. 3E, F), as described previously (F. Chen et al., 2015). Measurement lengths between pairs of points after expansion were compared to the distance expected given uniform expansion, and the average fractional deviation plotted as a function of measurement length (Fig. 3G). For a large acquisition of 42 fields of view (~650×750 μm after expansion), the measurement error was found to be a constant fraction (3.2% ± 1.7) of the measurement length (Fig. 3G). We used the similarity transform to calculate the overall expansion factor of the entire imaged area, and found it to be 9.4x, consistent with the expansion expected for the whole gel. Together, these data show that TREx enables uniform single step, ten-fold expansion that retains nanoscopic detail over large distances, in both cultured cells and thick tissue slices, with equal or better performance compared with the original 4x ExM.

### TREx enables subcellular localization of proteins and cellular ultrastructure in cultured cells

We next explored the use of TREx for high-resolution imaging of specific proteins, NHS stains and lipid membranes in cultured cells. For membranes, a custom-synthesized membrane probe compatible with the ExM process has been shown to visualize membranes in fixed brain tissue (Karagiannis et al., 2019). This probe relies on a peptide-modified lipid tail that intercalates in target membranes and provides opportunities for anchoring to the gel through D-lysines in its peptide sequence. We asked whether the commercially available membrane-binding fluorophore-cysteine-lysine-palmitoyl group (mCLING) could also be used for membrane staining and gel anchoring. mCLING has been developed as a fixable endocytosis marker consisting of a fluorophore and a short polypeptide group with one cysteine and seven lysines coupled to a palmitoyl membrane anchor (Revelo & Rizzoli, 2016). Due to the presence of multiple lysines, we hypothesized that mCLING would be compatible with standard ExM anchoring through AcX. While the standard protocol for mCLING delivery relies on active endocytosis in living cells, we tested whether mCLING would stain intracellular membranes more uniformly when added to fixed cells, which would have the added benefit of not perturbing intracellular membrane trafficking by long incubation in live cells. To test this, we fixed activated Jurkat T cells, incubated the fixed cells with mCLING overnight, and proceeded with the TREx protocol. We found that mCLING efficiently intercalates in both the plasma membrane and internal organelles and is retained following our standard anchoring procedure (Fig. 4A and Fig. 4—supp. Movie 1).

**FIGURE 4:**
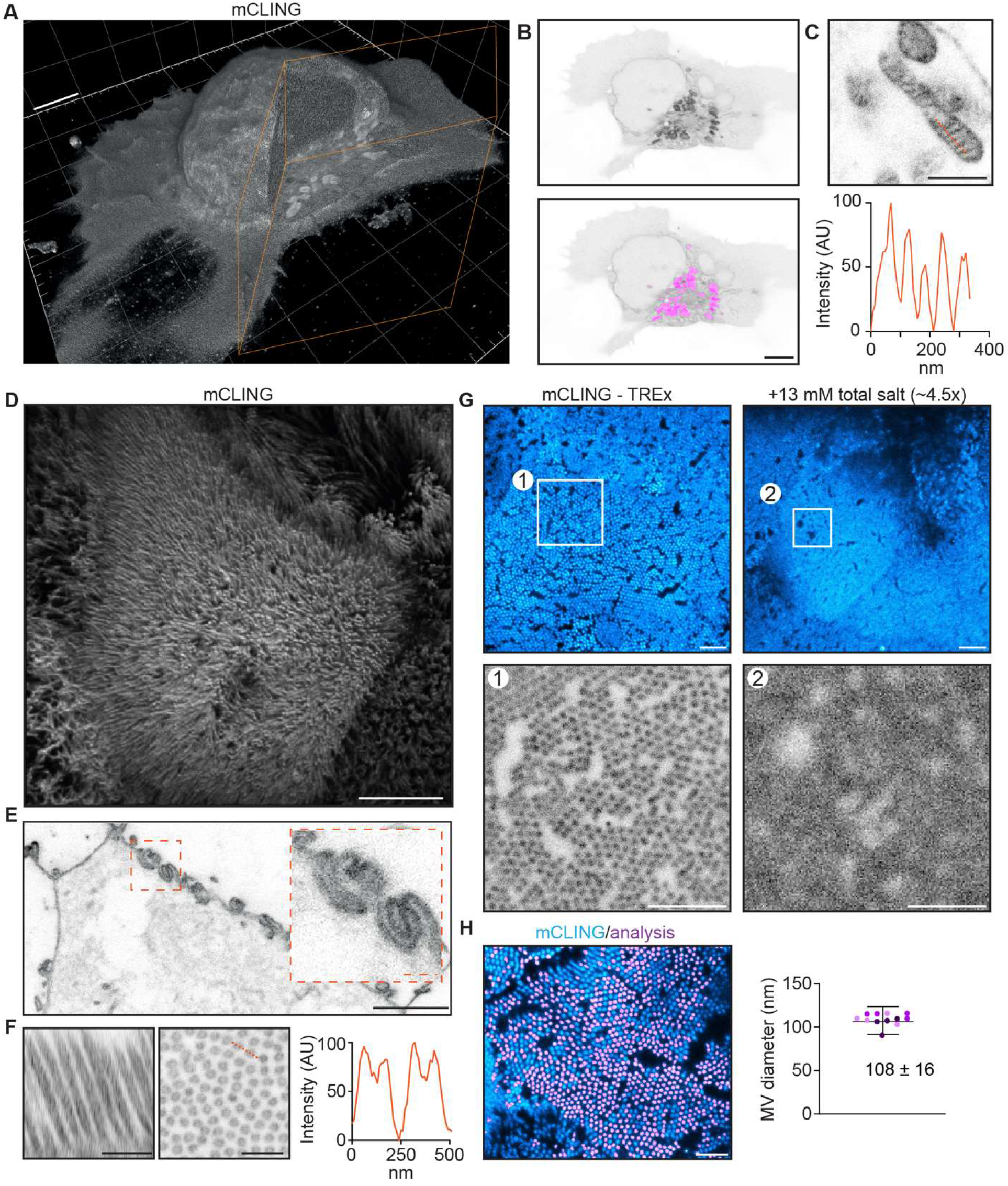
TREx can be used to visualize the ultrastructure of cellular membranes. A) Volumetric render of Jurkat T cell activated on anti-CD3 coated coverslip fixed and stained using mCLING. Colored clipping planes indicate portion clipped out to reveal intracellular detail. B) Immunological synapse of activated T cell in (A) revealing organelle clustering at the immunological synapse. Below: mitochondria segmented using the trainable Weka segmentation algorithm indicated in magenta. C) Representative example of mitochondrion in T cells visualized with mCLING. Line profile along the orange dashed line indicates mitochondrial cristae. D) Depth-coded volumetric projection of Caco2 monolayer apical brush border as seen from above looking down on the cells. E) Representative plane below the apical surface revealing highly interdigitated cell-cell contacts. F) Resliced (left) representative zoom (right) of brush border showing microvilli as hollow protrusions. Linescan indicated in orange. G) Comparison of dense brush borders after tenfold expansion in water (left) and ~4.5 times expansion in 13 mM salt (right, see Fig. 4—supp. fig. 1). Single plane of brush border and plane of same cell below the apical surface shown in cyan. Zooms 1, 2 correspond to areas of the same size corrected for the expansion factor to illustrate the increase in resolution of tenfold expansion. H) Quantification of microvilli diameter by determining the area of cross-sectioned (left). Plotted mean ± S.D. (107.7 ± 16.1 nm) of 12,339 microvili with means of individual cells color coded per replicate overlayed (4 cells per replicate, N=3). Scale bars (corrected to indicate pre-expansion dimensions): A, B, D, E (main) ~2 μm, C, E (zoom), F ~500 nm, G and H ~1 um.

By carefully rendering the imaged volumes we could, with one probe, both appreciate the ruffled morphology of the plasma membrane on top of the flattened part of the cell and visualize the organelle clustering typical of activated T-cells (Fig. 4A, B). As in electron microscopy, where distinct morphologies are used to identify organelles, we could clearly identify different organelles based on mCLING, suggesting that it could be used for automated segmentation of organelles. Indeed, we found that mitochondria could be readily segmented using a trainable Weka segmentation algorithm (Fig. 4B) (Arganda-Carreras et al., 2017). While the resolution of subcellular structures is limited by the density of mCLING moieties in the membrane, the efficiency of crosslinking to the gel, and the maximum expansion factor, we found TREx allows sufficient single-step expansion to resolve individual mitochondrial cristae (Fig. 4C), which are known to be as closely spaced as 70 nm (Stephan, Roesch, Riedel, & Jakobs, 2019). Although mCLING is membrane impermeable in live cells (due to multiple positively charged amino groups), it readily stained fixed and unpermeabilized cells following extended incubation. Because this approach does not require labeling live cells and is expected to reduce differences in uptake efficiency between intracellular compartments, we used this approach in all subsequent experiments.

We next tested if mCLING could also be used to visualize membranes in more complex cell types. To test this, we used differentiated Caco-2 cells grown to form an epithelial monolayer. Using TREx, we could expand the entire monolayer and visualize membranes using mCLING (Fig. 4D-H and Fig. 4—supp. Movie 2). The advantage of optical, volumetric imaging is underscored by the fact that we can easily render one dataset in several ways, either resembling scanning electron microscopy to highlight volumetric surface morphology (Fig. 4D), or transmission electron microscopy to explore single planes in more detail (Fig. 4E, F). For example, we were able to resolve the elaborate interdigitated cell-cell junctions that could previously only be clearly appreciated using electron microscopy (Drenckhahn & Franz, 1986), as well as resolve individual microvilli as hollow membrane protrusions within the dense brush border. To underscore the significant resolution increase of TREx compared to standard ExM we incubated expanded TREx gels with solutions of increasing ionic strength to shrink the gel back to ~ 4.5 times the size of the pre-expanded gel (Fig. 4G and Fig. 4—fig. sup. 1). When the 10x and 4.5x expanded gels were imaged, dense brush borders of differentiated cells could only be resolved in the 10x gel (Fig. 4G). To validate the expansion factor, we quantified the diameter of individual microvilli, as these have been thoroughly characterized with EM with a diameter of ~100 nm (Scott W, Mark S, & Matthew J, 2014). Indeed, we found an average diameter of 1.08 ± 0.16 μm (n=12339 from 12 cells, N=3), which corrected for an expansion factor of 10 is within the expected range. Together, these data illustrate the robustness of TREx in expanding multiple cell types and show how the increased expansion factor combined with a commercially available membrane stain provides rapid volumetric insights into the elaborate membranous architecture of cells.

Previously, we have used ExM to study the three-dimensional organization of microtubules (MT) in neurons and T cells (Hooikaas et al., 2020; Jurriens, van Batenburg, Katrukha, & Kapitein, 2020; Katrukha, Jurriens, Pastene, & Kapitein, 2021). For high-resolution imaging of the MT cytoskeleton, cells are usually pre-extracted with detergent and glutaraldehyde to remove the soluble pool of tubulin, followed by paraformaldehyde fixation (Tas et al., 2017). This reduces background but does not preserve membranes. We reasoned that the increased expansion of TREx would dilute the soluble tubulin background by the expansion factor cubed. Ten-fold expanded microtubules remain diffraction limited in width (i.e., 250-350 nm), so their signal should be reduced only by the expansion factor itself (due to the expansion along their length). Therefore, we asked whether the resulting relative boost in signal over background would eliminate the need for pre-extraction to enable high-resolution imaging of microtubules in combination with membranes. To test this, we fixed cells without pre-extraction, treated them with mCLING, stained for tubulin, and imaged the stained cells both before and after expansion with TREx (Fig. 5A, second panel). Expanded cells retained high quality anti-tubulin antibody signal exhibiting high contrast relative to the cytosolic background. We also observed increased detail in both mCLING and tubulin stains after expansion compared to before expansion, which was particularly apparent with side views of the same cell (Fig. 5A, far right). We next fixed cells expressing GFP-Sec61β without pre-extraction, treated them with mCLING, stained for GFP and tubulin, and then proceeded with TREx (Fig. 5B and Fig. 5-supp. Movie 1). Subsequent confocal microscopy revealed the interplay between microtubules and ER in three dimensions and revealed how other membranous organelles were connected to both structures (Fig. 5B, bottom). Thus, TREx facilitates high-resolution three-dimensional mapping of specific cytoskeletal and membranous structures in combination with markers that provide ultrastructural context.

**FIGURE 5:**
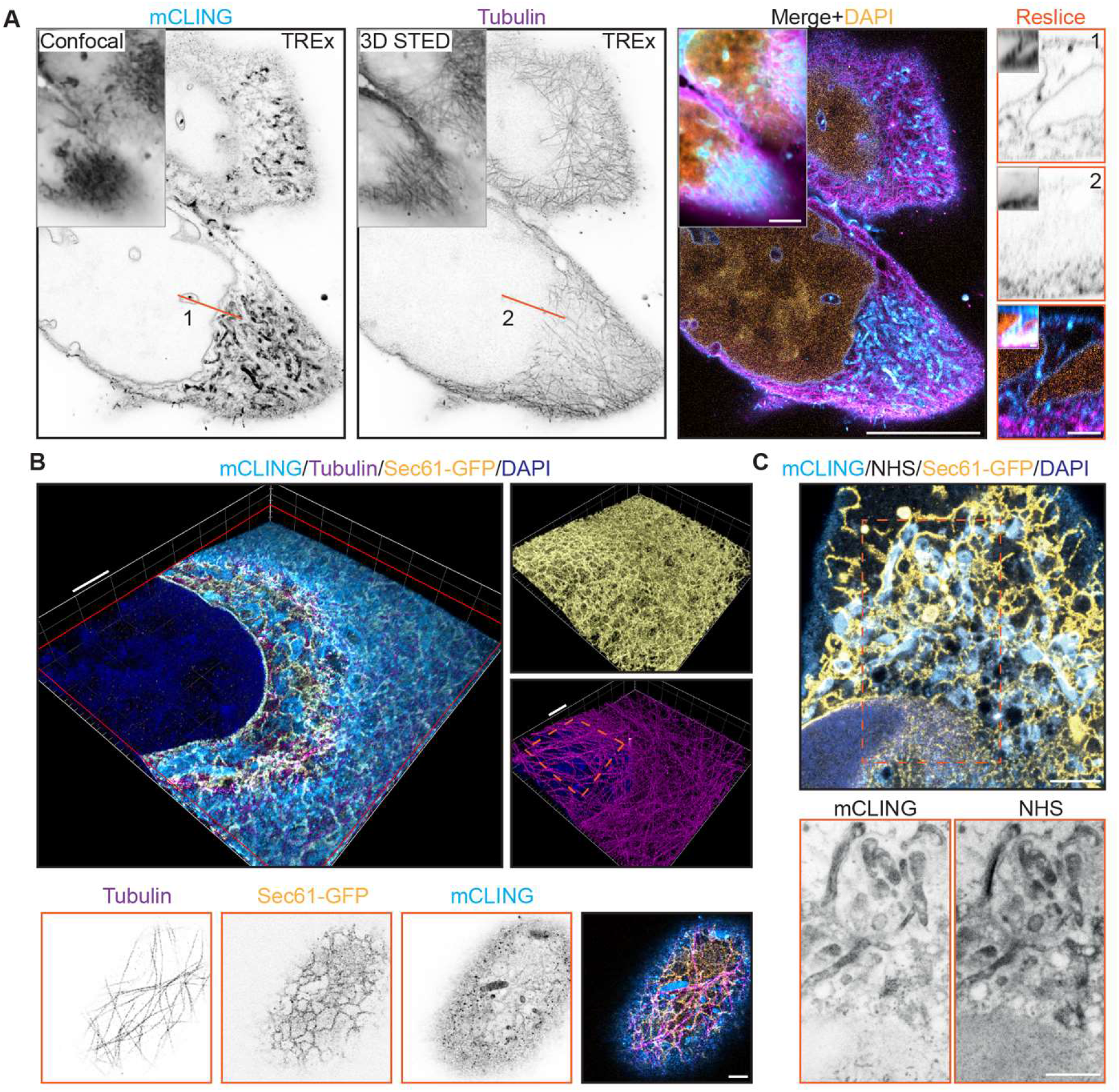
TREx Microscopy can combine antibody-based staining with NHS ester total protein stain to provide subcellular context. A) Single and merged planes of expanded U2OS cell stained for mCLING, tubulin and DAPI, grey outlined inserts show similar confocal and 3D STED acquisitions pre-expansion, for mCLING and tubulin respectively. Single planes of mCLING and tubulin are displayed in inverted contrast. Orange line (1,2) correspond to reslices (left) with inserts showing similar resliced planes pre-expansion. B) Volumetric render of U2OS cell expressing GFP-Sec61β stained for mCLING, GFP and tubulin. Top portion of cell is clipped with clipping plane indicated in red. Volumetric render of entire volume for GFP and tubulin in insert A and B, respectively. Single planes displayed in inverted contrast of the top of cell in B revealing the tight spatial organization below. C) Merged plane of expanded U2OS cell expressing GFP-Sec61β stained for mCLING, GFP, NHS ester and DAPI. Single planes of mCLING and NHS ester are displayed in inverted contrast. Scale bars (corrected to indicate pre-expansion dimensions): A (main) ~5 μm, B (renders) ~2 μm, A (reslices), B (single planes, below), C ~1 μm.

Finally, we tested whether TREx using general membrane stains could be combined with general protein stains and/or antibody stains. U2OS cells transfected with GFP-Sec61β were fixed, treated with mCLING, stained for GFP, and expanded with TREx followed by the NHS stain (Fig. 5C). Because we performed the NHS stain after disruption, we used high-temperature denaturing disruption, rather than proteolytic digestion. We found that secondary antibodies that had been used to stain GFP before gelation withstood this disruption step. We observed a clear degree of overlap between mCLING and NHS, especially in the dense perinuclear region, but we could also identify distinct features of each stain (Fig. 5C, bottom). These results demonstrate that general stains for membranes and proteins can be combined with antibody-based labeling to reveal specific proteins in their ultrastructural context.

## DISCUSSION

We developed Ten-fold Robust Expansion (TREx) in order to expand biological specimens 10-fold in a single round of expansion, without specialized equipment or procedures. In developing this method, we established a framework for assessing gel recipes operating near this apparent limit of single-round expansion. We found that the mechanical performance of gel recipes, i.e. resistance to deformation versus gel expansion factor, varies smoothly with changes in crosslinker. For all high monomer (~3 M total acrylamide and acrylate) gel recipe families, the relation between expansion factor and crosslinker concentration fell close to a common curve. The high radical initiation rate of family B enabled gelation without the inclusion of a crosslinker, suggesting that side reactions and polymer entanglement in these conditions create sufficient network crosslinks to form a gel. Gel deformation measurements plotted versus expansion factor, though less precise, also show high-monomer recipe families falling close to a common curve. Compared to the high-monomer families, the original ExM recipe family is less resistant to deformation for a given expansion factor and expands more for a given crosslinker concentration. The factor determining gel properties is not crosslinker concentration in the gel recipe *per se*, but rather the density of effective crosslinks formed between neighboring polymer chains in the gel. This suggests that the original, low-monomer recipe less efficiently incorporates crosslinker molecules as network crosslinks. This may be because the resulting lower rate of chain extension allows incorporated crosslinker molecules to be rereacted by the same polymer chain before they can react with neighboring polymers.

While the expansion factor of the original ExM recipe can be tuned by varying the crosslinker concentration, it has been shown that increasing the monomer content is required to maintain nanoscale isotropy, using centrioles as a convenient standard reference structure (Gambarotto et al., 2019). Considering gel quality versus expansion factor alone, the high monomer recipe family B derived from the U-ExM recipe allows for a 10-fold expanding gel (at crosslinker concentration of 30 ppm) with a low deformation index of 0.13. However, the high radical initiation rate used in this family (5 ppt APS and TEMED) results in fast gelation. This increases the time sensitivity of mounting the specimen in the gelation chamber and adds an additional challenge for adapting the method to thick tissues, requiring extended incubation in the gelation solution. Recipe families C and D solve this problem by reducing initiation rates, at a slight expense of mechanical performance compared with family B. Like family B, family C has a high acrylate content, which might contribute to imaging artifacts due to shrinkage of the sample prior to gelation, and inconsistent acrylate purity. Family D reduces the acrylate content by half while retaining similar mechanical performance to family C, especially in the ten-fold expansion regime. Finally, with family E we explored whether increasing the gelation temperature to 50 °C would produce the improved mechanical performance of family B (through increased temperature-dependent radical initiation) without premature gelation at room temperature. However, we found that this reduced the expansion factor and increased susceptibility to experimental variation. Therefore, we based our TREx recipe on recipe family D. The exact crosslinker concentration that produces 10-fold expanding gels was found to vary between labs (i.e. 50 ppm in Ashburn, VA, USA versus 90 ppm in Utrecht, The Netherlands, possibly due to differences in gelation chamber design), so we recommend that each lab test a range of crosslinker concentrations between 30 and 100ppm using their choice of specimen, gelation chamber, and incubator.

Earlier work has used the well-known architecture of the nuclear pore complex to compare macroscopic and nanoscopic expansion factors. For the conventional 4x expansion approach this revealed that the NPC diameter was 14-29% smaller than expected (Pesce et al., 2019; Thevathasan et al., 2019), suggesting that protein-dense complexes may resist full expansion. Using TREx, we found a NPC diameter that was only 8% smaller than the expected value. Further optimization of anchoring and disruption conditions may improve expansion uniformity for protein-dense structures such as NPCs. For applications requiring precise measurements, this may need to be validated for different structures individually. We characterized the overall expansion isotropy by comparing microtubules before and after expansion, finding expansion-induced measurement errors on average 3.2% of a given measurement length. This is in line with previous expansion methods (F. Chen et al., 2015; Tillberg et al., 2016) and is not a limiting factor for most biological applications.

We applied TREx to mouse brain tissue slices stained either for specific targets with antibodies, or for total protein distribution with NHS ester dyes. Single round ten-fold expansion with TREx followed by total protein staining was sufficient to reveal densely packed axons and dendrites running through the neuropil, while individual organelles could be resolved within the neuronal soma. The nuclear envelope, along with presumptive nuclear pore complexes, was also clearly resolved. The correct relative localizations of pre- and post-synaptic markers and pre-synaptic neurotransmitter vesicles stained with standard immunofluorescence were also retained in TREx-expanded tissue slices. For this characterization, we used a hybrid disruption approach incorporating reduced proteinase K digestion followed by high-temperature denaturation. We adopted this hybrid approach because we have noticed that sometimes the reduced protease treatment alone produces under-expanded nuclei while the protease-free treatment alone produces under-expanded synapses. In general, a higher degree of anchoring requires higher disruption strength to achieve full expansion on the macroscopic level. Further application-specific optimization may be beneficial, given the heterogeneity of biological tissue. For applications where maximizing total protein retention is not a priority, we recommend simply using a high concentration of proteinase K (e.g. 1:100 dilution, overnight).

We further demonstrated the utility of TREx for the study of cell biology through combinations with several staining modalities in cultured cells, prepared in several culture formats. After a single round of expansion with TREx, the commercially available membrane stain mCLING was able to clearly resolve the internal structure of mitochondria and the detailed pattern of plasma membrane ruffling in activated Jurkat T cells. While these structures would be readily resolved with electron microscopy of mechanically sectioned cells, we were able to do so in the context of complete cells, enabling concomitant detection and automated segmentation of mitochondria clustered followed T cell activation. Caco-2 cells grown on permeable filters were also successfully stained with mCLING and expanded with TREx to reveal the detailed structure of epithelial microvilli and membrane interdigitations at the contacts of neighboring cells. These structures had previously been known from electron microscopy but were now imaged with ease in the context of entire cell monolayers. Successful application of TREx to filter-cultured Caco-2 cells further demonstrates the robustness of TREx, because we had repeatedly failed to cleanly recover epithelial cultures using standard ExM. The robustness of TREx has been further demonstrated by its adoption in other biological systems (Gros, Damstra, Kapitein, Akhmanova, & Berger, 2021) including in cultured neurons (Özkan et al., 2021) and primary cultured human cells (Nijenhuis et al., 2021), and by its superior mechanical properties as measured by traditional materials characterization methods (R. Chen et al., 2021).

Combining mCLING with a total protein stain using NHS ester dye and an antibody marking the endoplasmic reticulum (ER) in U2OS cells, shows close contacts between ER and other organelles. While the NHS ester and mCLING staining patterns were similar in their overall contours, some clear differences in staining patterns were noted, including the presence of presumptive nuclear pore complexes in the NHS ester channel. The strong overlap between NHS ester and mCLING stains was not unexpected, given the reactivity of NHS esters towards both unreacted lysines in the mCLING molecule and antibodies. However, the extent to which NHS ester staining is truly unbiased over all proteins and how it may be modulated by the local environment awaits further exploration.

In U2OS cells, TREx retained anti-tubulin antibody stain with high efficiency, maintaining continuous microtubules with high signal-to-noise ratio after 1000-fold volumetric expansion. Combined staining for ER and all membranes revealed close appositions along microtubules, ER and presumptive mitochondria. In unexpanded cells, high quality antibody staining for microtubules requires non-polymerized tubulin to be removed with a pre-extraction step, which destroys membranes and extracts other proteins. For specimens expanded with TREx, this is not necessary as the monomeric tubulin signal is diluted 1000-fold volumetrically, while the diameter of an expanded microtubule is still below the diffraction limit, so the signal is only diluted by the 10-fold linear expansion factor. This 1000-fold volumetric dilution is also what enables the use of dense protein stains such as NHS ester dyes, which in unexpanded specimens would be too dense to resolve any meaningful structure.

In summary, by systematically exploring the ExM recipe space we established a novel recipe using standard ExM reagents that has been rapidly adopted by other labs. TREx allows for ten-fold expansion of both thick tissue slices and cells in a single expansion step and has applications in tissue and high-resolution subcellular imaging. Importantly, TREx of antibody-stained samples can be combined with off-the-shelf small molecule stains for both membranes and total protein to localize specific proteins in their ultrastructural context.

## ACKNOWLEGDMENTS

We are grateful to Sven van IJzendoorn (UMCG) and Wilco Nijenhuis (UU) for providing the Caco2 monolayer samples. We thank the Janelia light microscopy core facility for the use of confocal and lightsheet microscopes. We thank the Janelia cell culture core facility for maintaining and providing cultured cells. We thank the Janelia histology core facility for providing tissue slices. A.A. is supported by the Netherlands Organization for Scientific Research Spinoza Prize. L.C.K. is supported by the European Research Council (ERC Consolidator Grant 819219). B.M., M.E., and P.W.T. are supported by the Howard Hughes Medical Institute (HHMI).

## MATERIALS AND METHODS

### Recipe space exploration

#### Gelation chambers

A glass slide served as the bottom piece of each gelation chamber. Four strips of 250 μm-thick adhesive silicone material (Digikey cat# L37-3F-320-320-0.25-1A), ~3 mm wide and running the width of the slide, were adhered to the slide to partition it into three separate chambers, each ~12 mm wide. A plus-charged glass slide was placed over the silicone strips to form the top of the gelation chamber and held in place with tape. Two sides of each chamber were open to air, providing a convenient fill port for adding ~100 uL of monomer solution after chamber construction.

#### Gel synthesis and characterization

Sodium acrylate was made by neutralizing acrylic acid (Sigma, 147230) with NaOH until the pH reached the range of 7.5-8. Initial neutralization (until pH ~7) was done with 10 N NaOH on ice and using a fume hood. Neutralization was done in a volume of water calculated to yield a final concentration of 4 M sodium acrylate. The gel recipes for each family contained 1x PBS and the amounts of acrylamide (Sigma, A4058), sodium acrylate, and initiator (APS, Sigma, A3678) indicated in Fig. 1A. Each gel recipe contained the same amount of TEMED (Sigma, T7024) as APS. For each recipe family, gelation solution with crosslinker withheld (but including APS and TEMED) was premixed on ice, in one tube for each recipe family. This solution was then split into six tubes and mixed with serial dilutions of crosslinker (bisacrylamide, Sigma, M1533) to yield complete gelation solution with final crosslinker concentrations (in ppm) of 1000, 300, 100, 30, 10, and 0. Complete gelation solution was pipetted into gelation chambers and incubated at 50 °C for 1 hour (family E) or 37 °C for 2 hours (families A-D). Gels were then cooled for 15 minutes at room temperature and chamber tops carefully removed. Gels typically remained stuck exclusively to the top (plus charged) slide. Samples of each gel were taken with a 6 mm biopsy punch, taking care to avoid material within ~2 mm of the chamber edges (to avoid oxygen exposure from air or silicone material during gelation). Excess gel was scraped away with a razor blade. A few drops of distilled water were pipetted onto each gel to help release them from the glass slide. Each 6 mm gel specimen was gently released from the slide with a razor blade, placed in a 9 cm Petri Dish and expanded by washing with excess water 2x 15 minutes followed by 2x 1 hour. Diameters of expanded gels were measured and divided by 6 mm to obtain the expansion factor. A semi-circle 25 mm in diameter was punched from each gel using a cookie cutter. Semi-circular gel punches were placed in a plastic tray, which was stood up on end so that the gel stood upright on its curved side, allowing the flat edge to deform under the force of gravity. Each gel was photographed, with a ruler positioned for scale. Using ImageJ, each top edge was described by seven manually chosen points, which were then fit to a circle. This best-fit circle was used to calculate the vertical deviation of the gel corners, which was divided by the gel radius to obtain the deformation index.

#### TREx gelation solution

Sodium acrylate was either purchased (Sigma, 408220) or made by neutralizing acrylic acid as described above. TREx gelation solution contains 1.1 M sodium acrylate, 2.0 M acrylamide (AA), 50 ppm (for tissue slices and cultured cells prepared at Janelia) or 90 ppm (for cultured cells prepared at Utrecht University) N,N’-methylenebisacrylamide (bis), PBS (1x), 1.5 ppt APS, 1.5 ppt TEMED, and (optionally, for thick tissue slices) 15 ppm 4-hydroxy TEMPO (4HT, Sigma, 176141). Monomer solution was made by combining all components of gelation solution except APS, TEMED, and 4HT. Monomer solution may be aliquoted and stored at −20°C, but must be thawed at room temperature and vortexed before use to redissolve any acrylamide crystals that may have precipitated at low temperature before freezing. Fully dissolved monomer solution may be kept on ice for up to several hours before crystallization will occur. 4HT, TEMED, and APS were added to monomer solution to produce gelation solution directly before use.

### Tissue experiments

#### Fixation and antibody staining

Mice were transcardially perfused with ice cold 4% formaldehyde in 100 mM sodium phosphate buffer, pH 7.4. Brains were dissected out and post-fixed in 4% formaldehyde at 4°C overnight (Fig. 2A) or for 2 hours (Fig. 2B), followed by washing with PBS (1x) and slicing by vibratome at 100 μm. For Fig. 2B, slices were stained with standard IHC procedures. Primary antibodies were used at 1:300 dilution in PBS with 0.1% Triton and 2% BSA (PBT) overnight at 4°C (Chicken anti-Bassoon Synaptic systems cat#141016, RRID:AB_2661779; Rabbit anti-Homer abcam cat#97593, RRID:AB_10681160; Mouse IgG3 anti-VGAT SySy cat#131011, RRID:AB_887872). Sections were washed 3x 30min in PBT and stained for at least 6 hours in secondary antibodies 1:500 in PBT at room temperature (Goat anti-Rabbit Alexa 488 Abcam cat#150077, RRID:AB_2630356; Goat anti-Mouse IgG3 Alexa 594 Invitrogen cat# A-21155, RRID: AB_2535785; Goat anti-chicken CF633 Biotium cat#20126, RRID: AB_10852831). Stained sections were washed 3x 30min in PBS.

#### TREx

Brain slices were treated with 100 μg/mL (Fig. 2A) or 10 μg/mL (Fig. 2B) acryloyl-X SE (ThermoFisher, A20770) in PBS (diluted from a 10 mg/mL anhydrous DMSO stock solution) for 1 hour at room temperature, followed by rinsing with PBS. Slices were incubated with TREx gelation solution (using 50 ppm bis, and with 4HT added up to 15 ppm), for 20 min on ice, with shaking. Following incubation on ice, each tissue specimen was placed on a glass slide at room temperature. Four dabs of vacuum grease were applied to the slide, with each dab at least several mm from the tissue specimen. A coverslip was placed over the tissue and vacuum grease dabs, and pressed down until contacting the tissue, taking care not to let the tissue slide around on the slide. The vacuum grease served to hold the assembly in place, thus forming the gelation chamber. Gelation solution was pipetted into the chamber from the side to fully surround the tissue. The chamber was incubated at 37 °C for 1 hour to complete gelation. Following embedding, excess gel was removed with a razor blade, and gelled slices were recovered into PBS. The gel for Fig. 2B was digested in proteinase K (NEB, P8107S) diluted 1: 1000 in PBS for 3 hours at room temperature and washed in PBS 4x 30 min. Gels for both Fig. 2A, B were then placed into disruption buffer (5% SDS, 200 mM NaCl, 50 mM Tris pH 7.5) in a 2 mL Eppendorf tube and incubated at 80 °C for 3 hours followed by rinsing in 0.4 M NaCl and washing 2x 30 min in PBS. Gels were stained with BODIPY-FL NHS (total protein stain) at 10-20 μM (Fig. 2A) or DAPI at 200 μg/L (Fig. 2B) in PBS for 1 hour at room temperature. Gels were placed in glass bottom 6-well plates and washed in milliQ water 3x 15 minutes followed by 2x 1 hour to fully expand. Gels were imaged using a Zeiss LSM 800 confocal microscope with 40x/1.1NA, water immersion objective (Fig. 2A) or Zeiss Z1 lightsheet microscope with 10x/0.3NA illumination objectives and 20x/1.0NA water immersion detection objective (Fig. 2B).

#### Image processing

For Fig. 2A, raw data was drift corrected using Huygens Professional (SVI) and imported into ImageJ where a sum-projection of 2 planes (z-spacing: 0.8 μm) was made. Fig. 2B is a maximum projection of 2 planes (z-spacing: 0.38 μm) and indicated zoom is a volumetric render of the raw data in Arivis.

#### Synaptic distance

Raw data was segmented using ilastik Pixel and Object segmentation workflows (Berg et al., 2019). For each Homer-positive segmented object (post-synaptic compartment), the closest Bassoon-positive segmented object (pre-synaptic compartment) was selected. Synaptic distance was defined as the distance between the local peaks in intensity that were closest to the mask center of mass.

### Nuclear Pore Complex experiment

#### Cell culture, fixation, and antibody staining

U2OS cells with homozygous GFP-NUP96 knock-in (Cell Lines Service, no. 195) were maintained in DMEM (Corning) supplemented with 10% FBS (Gibco), 1% L-glutamine (Gibco), and 1% penicillin-streptomycin (Gibco). Exponentially growing cells were harvested and seeded onto 12 mm, No. 1 coverslips (Carolina Biological Supply) for use in Expansion Microscopy. Cells were grown at 37 °C and 5% CO_2_. Cells were fixed with 4% formaldehyde (EMS, RT 15714) in 1x PBS for 10 minutes at room temperature, then rinsed with 1xPBS. Cells were stained with standard immunocytochemistry (ICC) procedures. Primary antibodies were used at 1:200 dilution in PBS with 0.1% Triton and 2% BSA (PBT) for 2 hours at room temperature (Chicken anti-GFP, Aves cat#GFP-1020, RRID:AB_10000240; Rabbit anti-NUP153, Abcam cat#ab84872, RRID:AB_1859766), followed by washing 3x 5 min in 1x PBS. Secondary antibodies were used at 1:200 dilution in PBT for 2 hours at room temperature or at 4 °C overnight (Goat anti-Chicken Alexa 488, ThermoFisher Scientific cat#A11039, RRID_AB2534096; Goat anti-Rabbit Alexa 594, ThermoFisher Scientific cat#A11037, RRID_AB2534095), followed by washing 3x 5 min in 1x PBS. Stained cells were imaged before expansion on an epifluorescence microscope, Nikon Ti-E with 60x/NA1.2 water immersion objective. The imaged region was indicated by marking the back of the coverslip with a marker.

#### TREx

Fixed cells were anchored with 100 μg/mL AcX in 1xPBS for 1 hour at room temperature and embedded using the TREx gelation solution. The gelation chamber was constructed from a 20 mm diameter, adhesive-backed silicone gasket (Sigma, GBL665504) affixed to a glass slide. The 12 mm coverslip with cultured cells was affixed to the center of the gelation chamber with a dab of vacuum grease and covered with PBS. TEMED and APS were then added to the TREx monomer solution on ice and mixed well to produce gelation solution. The PBS was tipped off from the cells, which were rinsed with ~100 μL of gelation solution. ~200 μL of gelation solution was placed into the gelation chamber, which was sealed with a 22 mm-square #2 coverslip. The completed gelation chamber was placed at 37 °C for 1 hour to complete gelation. The chamber was disassembled, and the gel carefully trimmed with a curved scalpel into a right trapezoid shape centered around the pre-gelation imaged area. The trimmed trapezoid was photographed with a ruler for scale, quickly to avoid shrinking due to evaporation, and recovered into PBS. The gel was then digested with proteinase K (NEB, P8107S) diluted 1:1000 in PBS for 3 hours at room temperature and washed in PBS 4x 30 min. Digested gels were placed into disruption buffer (5% SDS, 200 mM NaCl, 50 mM Tris pH 7.5) in a 2 mL Eppendorf tube and incubated at 80 °C for 3 hours followed by rinsing in 0.4 M NaCl and washing 2x 30 min in PBS. Disrupted gels were expanded fully with several washes in deionized water, photographed again with a ruler for scale, and imaged with a Zeiss LSM 800 confocal microscope with 40x/NA1.1 water immersion objective.

#### Data analysis

The gel size before and after expansion was measured from the gel photographs. The centers of 60 randomly chosen NPCs in three non-adjacent cells were identified manually and saved as an ROI list in ImageJ. The radial intensity distribution of each NPC was computed using the “Radial Profile Plot” plugin (https://imagei.nih.gov/ii/plugins/radial-profile.html) and saved as a csv. Radial intensity distributions were loaded into Matlab for further processing. A Gaussian distribution was fit to a window in the middle of each profile and the center of the Gaussian was taken as the radius of the corresponding NPC.

### Wild type, transfected, and T-cell experiments

#### Cell culture

Jurkat T cells (clone E6.1) were grown in RPMI 1640 medium w/ L-Glutamine (Lonza) supplemented with 9% Fetal Bovine Serum and 1% penicillin/streptomycin. For T cell activation, 18 mm #1.5 coverslips (Marienfeld, 107032) were coated with Poly-D-Lysine (Thermo Fisher Scientific, A3890401), washed with phosphate buffered saline (PBS) and incubated overnight at 4 °C with a mouse monoclonal anti-CD3 (clone UCHT1, StemCell Technologies, #60011) 10 μg/mL in PBS. Cells were spun down for 4 minutes at 1000 rpm and resuspended in fresh, prewarmed RPMI 1640 medium, after which cells were incubated on the coated coverslips for 3 minutes prior to fixation. U2OS cells were cultured in DMEM medium supplemented with 9% Fetal Bovine Serum and 1% penicillin/streptomycin. U2OS cells were transfected with GFP-Sec61β (Addgene, 15108) using FuGENE6 (Promega). Caco2-BBE cells (a gift from S.C.D. van IJzendoorn, University Medical Center Groningen, The Netherlands) were maintained in DMEM supplemented with 9% FBS, 50 μg/μl penicillin/streptomycin and 2 mM L-glutamine. For imaging, cells were seeded on 6.5 mm Transwell filters (3470; Corning) at a density of 1 x 10^5^/cm^2^ and cultured for 10-12 days to allow for spontaneous polarization and brush border formation.

#### Immunofluorescence, mCLING treatment, and antibody staining

For all experiments, cells were fixed for 10 minutes with pre-warmed (37 °C) 4% paraformaldehyde + 0.1% glutaraldehyde in PBS. For visualization of lipid membranes, cells were washed twice in PBS after fixation and incubated in 5 μM either mCLING-Atto647N (Synaptic Systems, 710 006AT1) or mCLING-Atto488 (Synaptic Systems, 710 006AT3) in PBS overnight at RT. The following day, cells were fixed a second time with pre-warmed (37 °C) 4% paraformaldehyde + 0.1% glutaraldehyde in PBS. Next, cells were washed with PBS and permeabilized using PBS + 0.2% Triton X-100. Epitope blocking and antibody labeling steps were performed in PBS + 3% BSA. For immunofluorescence staining, we used a rabbit monoclonal antibody against α-tubulin (clone EP1332Y, Abcam, ab52866) and a chicken polyclonal antibody against GFP (Aves Labs, GFP-1010) in combination with goat anti-rabbit IgG (H+L) Alexa Fluor 594 (Molecular Probes, a11037) and goat anti-chicken IgY (H+L) Alexa Fluor 488 (Molecular Probes, a11039), respectively.

#### TREx

For TREx, samples were treated with 100 μg/mL acryloyl-X SE (AcX) (Thermo Fisher, A20770) in PBS overnight at RT. TEMED and APS were added to monomer solution (1.5 ppt each) to produce gelation solution. 170 μL of gelation solution was transferred to a silicone gasket with inner diameter of 13 mm (Sigma-Aldrich, GBL664107) attached to a parafilm-covered glass slide, with the sample put cell-down on top to close off the gelation chamber. The sample was directly transferred to a 37 °C incubator for 1 hour to fully polymerize the gel. All gels excluding samples that were processed for subsequent NHS ester staining were transferred to a 12-well plate and digested with 7.5 U/mL Proteinase-K (Thermo Fisher, EO0491) in TAE buffer (containing 40 mM Tris, 20 mM acetic acid and 1 mM EDTA) supplemented with 0.5% Triton X-100, 0.8 M guanidine-HCl, and DAPI for 4 hours at 37 °C. The gel was transferred to a Petri dish, water was exchanged 2x 30 minutes and the sample was left in milliQ water to expand overnight.

For NHS-staining, gels were first treated in disruption buffer containing 200 mM SDS, 200 mM NaCl and 50 mM Tris pH 6.8 for 1.5 hours at 78°C. Gels were washed twice for 15 minutes in PBS and incubated with 20 μg/mL Atto 594 NHS ester (Sigma-Aldrich, 08471) in PBS prepared from a 20 mg/mL stock solution in DMSO for 1 hour at RT with shaking. After staining, gels were washed with excess of PBS, transferred to a Petri dish, and expanded overnight. Prior to imaging the cells were trimmed using a scalpel blade to fit in a Attofluor Cell Chamber (Molecular probes A-7816).

#### Image acquisition and analysis

ExM and pre-expansion images were acquired using a Leica TCS SP8 STED 3X microscope equipped with a HC PL APO 86x/1.20W motCORR STED (Leica 15506333) water objective. A pulsed white laser (80 MHz) was used for excitation, when using STED a 775 nm pulsed depletion laser was used. The internal Leica GaAsP HyD hybrid detectors were used with a time gate of 1 ≤ tg ≤ 6 ns. The set-up was controlled using LAS X.

All data processing and analysis was done using Matlab, ImageJ and Arivis.

Fig. 3D panels are maximum intensity projections of the bottom ~1 μm of cells. For Fig. 3E-F, BigWarp (Bogovic, Hanslovsky, Wong, & Saalfeld, 2016) was used to manually pick control points for non-rigid registration. The analysis scripts “bigwarpSimilarityPart.groovy” and “Apply_Bigwarp_Xfm_csvPts.groovy” were used to calculate deformation fields that register expanded images to pre-expansion images, and decompose each deformation field into a similarity part (corresponding to theoretical ideal expansion) and a residual elastic part (thin-plate spline, corresponding to non-ideal deformations introduced by expansion), adapted from (Jurriens et al., 2020). The similarity part was used to find the macroscopic expansion factor, while the residual elastic part was used to calculate the measurement error as follows. A Matlab script was used to calculate the measurement error for all pairs of points in the image as described in (F. Chen et al., 2015) by finding the magnitude of the difference between the displacement vectors for each pair of points in the residual elastic deformation field. These differences were binned according to the distance between points in the pre-expansion image. For each measurement length bin, the mean and standard deviation of measurement errors was calculated and plotted.

Fig. 4A raw data was imported in Arivis, a Discrete Gaussian Filter with smoothing radius of 2 was applied and this dataset was used for volumetric renders and clipping. Gamma was adjusted manually to increase visibility of plasma membrane ruffles and intracellular organelles in the same view. For Fig. 4B, the same raw dataset was imported in ImageJ and a sum-projection of 3 planes (z-spacing: 0.35 μm) around the plane of the immunological synapse was segmented for mitochondria using the trainable Weka segmentation plugin in ImageJ. Fig. 4C is a sum-projection of 3 planes (z-spacing 0.35 μm). The linescan in Fig. 4C was generated using ImageJ and processed using Graphpad Prism 8. For Fig. 4D, raw data was imported in Arivis, a Discrete Gaussian Filter with smoothing radius of 2 was applied and this dataset was volumetrically rendered with the opacity mapped to the z-axis. Fig. 4E is a sum projection of 5 slices (z-spacing 0.35 μm). Fig. 4F-H are sum projections of 3 planes (z-spacing: 0.35 μm) and respective zooms. For the MV diameter analysis in Fig. 4H, sum projections of 3 planes were thresholded (ImageJ, set to auto), watershed to split joining particles and the area determined using the analyze particles function in ImageJ which was converted to diameter as in (Julio, Merindano, Canals, & Ralló, 2008).

Fig. 5A panels are sum projections of 3 planes (z-spacing before expansion and after expansion 0.07 and 0.15 μm, respectively), reslices are sum projections (3 planes) of resliced data. For Fig 5B, raw data was imported into Arivis, a Discrete Gaussian Filter with smoothing radius of 2 was applied and this dataset was used for volumetric renders and clipping. Shown single planes are sum projections of 3 slices (z-spacing 0.35 μm) of the same raw data and was processed using ImageJ. Fig 5C is a maximum projection of 3 planes (z-spacing: 0.35 μm).

---

**FIGURE 2—FIGURE SUPPLEMENT 1:**
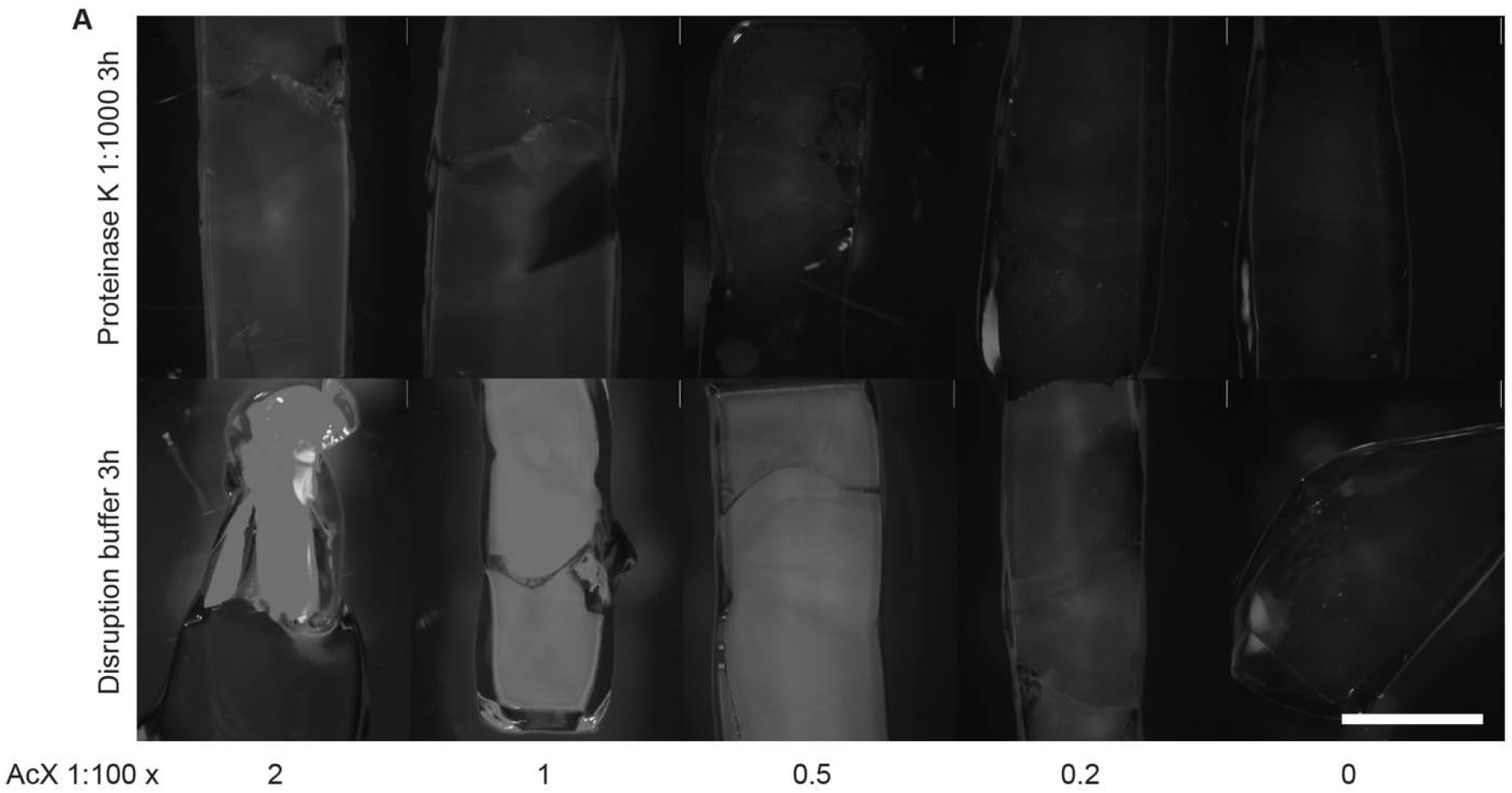
Comparison of anchoring and disruption conditions. A) Mouse brain tissue samples anchored with varying amounts of acryloyl-X SE (AcX), stained with NHS ester dye, and disrupted with two methods: proteinase K diluted 1:1000 into PBS and applied at room temperature (top row) or denaturing disruption buffer applied at 80 °C (bottom row) for 3 hours. AcX was diluted from a 10 g/L stock to 200, 100, 50, 20, and 0 mg/L (from left to right) into PBS and applied for 1 hour.

**FIGURE 4—FIGURE SUPPLEMENT 1:**
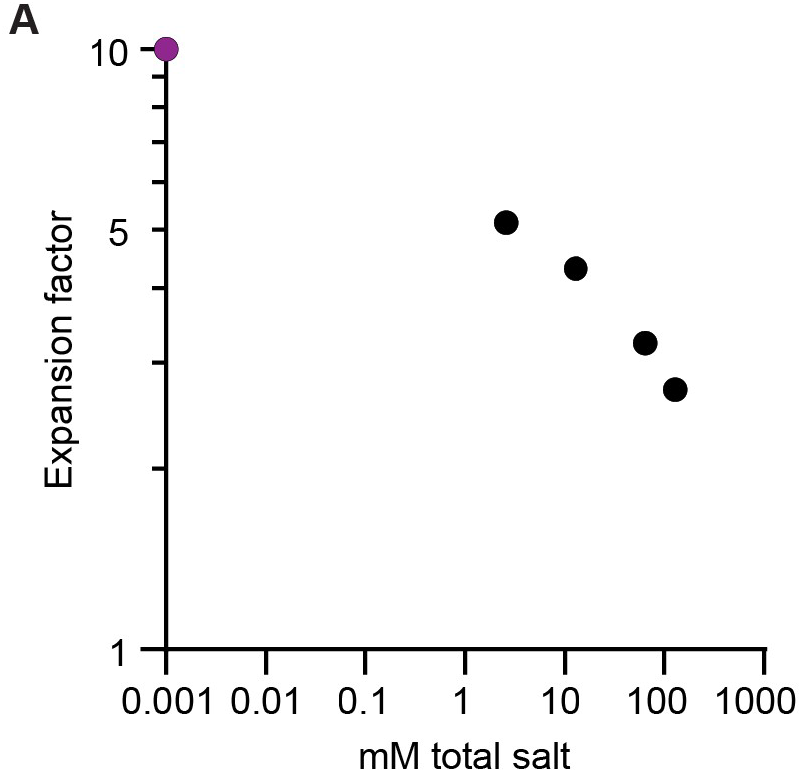
Expansion factor versus ionic strength. A) Expansion factor of TREx gel without biological sample as a function of ionic strength. Black dots are measured values with mM total salt being derived from dilutions of PBS (factor 1, 0.5, 0.1, 0.02 of PBS). The magenta dot represents 10x expansion in water in equilibrium with atmospheric CO_2_. Assuming each H^+^ corresponds to one HCO_3_^-^ ion, the measured pH of water in equilibrium with room air of 6 implies an ionic strength of 10^-6^, or 1 μM.

## SUPPLEMENTAL MOVIES

**FIGURE 2—Supplemental Movie 1:** Z-stack of Fig. 2A

**FIGURE 2—Supplemental Movie 2:** 3D render of Fig. 2B

**FIGURE 4—Supplemental Movie 1:** 3D render of Fig. 4A

**FIGURE 4—Supplemental Movie 2:** 3D render of Fig. 4D

**FIGURE 5—Supplemental Movie 1:** 3D render of Fig. 5B

